## Supplemental Figure S1 for "The molecular basis of Abelson kinase regulation by its αI-helix"

### Supporting Information

#### Supplementary Figures

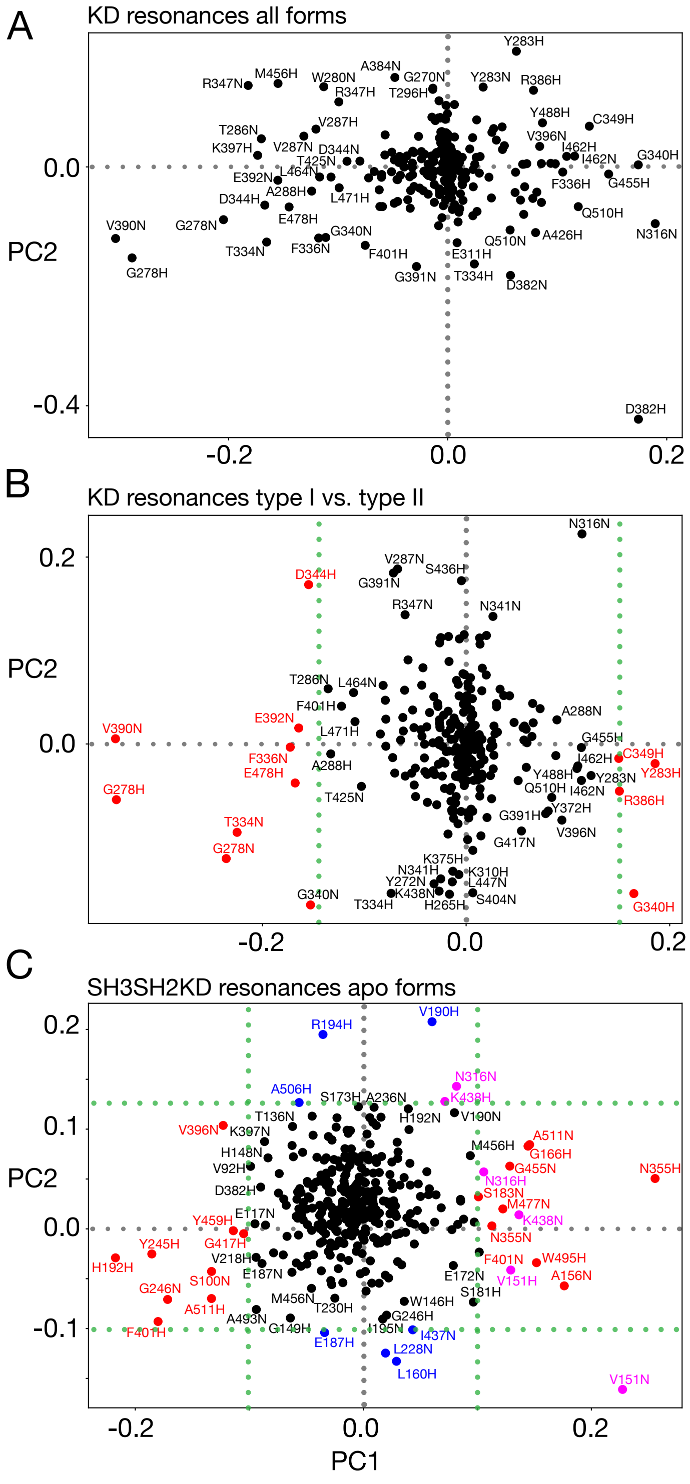


**Figure S1. PCA loadings showing the contributions of the $^{1}H$ and $^{15}N$ chemical shifts of individual residues to PC1 and PC2.** (**A**) PCA of all KD chemical shifts of all investigated complexes corresponding to Figure 4A. (**B**) PCA of KD chemical shifts for type I and type II inhibitor complexes (Figures 4B and 4C). Residues with |PC1| > 0.15 are indicated in red. (**C**) PCA of all SH3SH2KD shifts of all apo forms (Figure 4D and 4E). Residues with sizeable |PC1| > 0.1 or sizeable PC2 (< -0.1 or > 0.125) are indicated in red and blue, respectively. V151, N316, and K438 have both sizeable PC1 and PC2 and is shown in magenta.
